# Cross-kingdom regulation of tRNAs/tRFs derived from Chinese yew

**DOI:** 10.1101/821165

**Authors:** Kai-Yue Cao, Tong-Meng Yan, Ji-Zhou Zhang, Ting-Fung Chan, Jie Li, Chong Li, Elaine Lai-Han Leung, Zhi-Hong Jiang

## Abstract

Plants containing countless chemical constituent have benefited mankind since the origin of life. Although secondary metabolites in plants, such as morphine, artemisinin and taxol, have been developed as therapeutic drugs for clinical therapy, few study focuses on the pharmacological activities of plant small RNAs with function of cross-kingdom regulations. Yew is widely considered as a “superstar” plant due to the discovery of paclitaxel, or taxol, which is a well-known natural drug for the treatment of multiple types of cancer^1^. Here we show the surprising finding that an RNA fragment, named tRF-T11, derived from tRNA^His(GUG)^ of Chinese yew strongly suppressed human ovarian cancer progression. In A2780 cells, tRF-T11 mimic (a double-stranded RNA with tRF-T11 as antisense chain) exhibited potent cytotoxicity comparable to that of taxol, but no significant cytotoxicity to normal ovarian surface epithelial cells. Moreover, the cytotoxicity of tRF-T11 mimic is 80-fold stronger than that of taxol in taxol-resistant A2780 cells. Bioinformatic and molecular biological studies revealed that tRF-T11 targets transient receptor potential cation channel subfamily A member 1 (*TRPA1*) to inhibit its expression levels. In a further *in vivo* investigation, the growth rate of ovarian tumor xenografts in nude mice was significantly reduced by treatment with tRF-T11 mimic, and the *TRPA1* protein expression in tumors treated with tRF-T11 mimic was also down-regulated. Our findings are the first to provide evidence that plant-derived tRFs can regulate the expression of target genes *in vitro* and *in vivo*, indicating that they may become as a new source of druggable siRNA. Moreover, this discovery demonstrated a pilot example of an innovative approach for not only identifying pharmacologically-active tRFs from plants, but also for improving the efficiency and possibilities of discovering new drug target.

---

Plants have provided mankind with an important source of medicinal products for millennia. Throughout ancient history, humans learned to harness herbal decoctions for disease prevention and treatment. Today, thousands of secondary metabolites, such as flavonoids, terpenes and alkaloids, from this chemical treasure have been identified as active compounds. Intriguingly, crude plant extracts were more effective than the synthetic versions of phytochemicals, implying that whole plant extracts contain a cocktail of products in which substances like macromolecules other than small molecules may contribute to the overall pharmacological effects^2^. However, few study has been performed on the pharmacological activity of plant macromolecules excluding polysaccharides because they are considered inactive, nonabsorbable and unstable in human body, such as RNA, DNA and proteins. Small RNAs (<200 nt) were found to play critical roles in a wide range of biological processes in microorganisms, plants, animals and human, and to participate in a variety of human diseases by regulating protein-coding genes^3, 4^. Since the first evidence that plant-derived miRNAs can regulate gene expression in mammalian systems was provided by Zhang *et al*^5^ in 2012, many other miRNAs from small RNAs in plants have been extensively studied for their cross-kingdom regulatories^6–9^. These findings suggested the promise of small RNAs from plants to be developed as therapeutic agents. A recent report by Shekhawat^10^ showed that the small RNA of corn exhibited appreciable antiproliferative activity toward HeLa cells, implying the possibility that small RNA species with larger molecular weight than miRNA play regulatory roles. In fact, as the most abundant small RNA species expressed in all living organisms, transfer RNAs (tRNAs) play a well-defined role in protein translation and perform diverse regulatory functions in some important biological processes, such as cell signaling, apoptosis modulation and stress response programs^11^. Recently, tRNA-derived fragments (tRFs) have also been recognized to play regulatory roles^12^. This class of small RNAs is non-randomly generated from tRNAs, and many are specifically and abundantly produced under particular conditions, such as cancer and viral infection^13, 14^. Some studies have also revealed that the expression of several tRFs is positively correlated with cell proliferation in different cancer cell lines^15^, suggesting that tRFs and their related signaling pathways could be novel targets for intervention in the treatment of cancers. These findings prompted us to consider the further possibility that in addition to well-established players such as miRNAs and small molecules in plants, tRNAs and their fragments, which account for overwhelming majority of small RNA species in plants, may hold great potential as biological regulators in mammals. Below, we describe the first study of the pharmacological activities of tRNAs and tRFs derived from Chinese yew (*Taxus chinensis*) and identification of their molecular target.

Total RNA with high integrity (RIN=7.7) and purity (A260/280=2.08, A260/230=2.00; Supplementary Fig 1a) was isolated from fresh branches of *T. chinensis* by using a well-developed CTAB method (Supplementary Fig 1b, c), in which small RNA was separated from large RNA with high purity (<200 nt, Supplementary Fig 2). The tRNAs were then gel fractionated and purified (Supplementary Fig 3a) to successfully delete 5S and 5.8S rRNA and other RNA species (Supplementary Fig 3b). Sequence information and the proportion of each type of RNA in this tRNA-enriched fraction (tEF) were determined by performing next-generation sequencing (NGS). As shown in Fig 1c and Supplementary Table 1, over 1.7 million clean reads distributed mainly from 60 to 80 mer were generated, and over 20 individual tRNA sequences were identified, which accounted for approximately 14% of the overall number of reads in the tEF library (Fig 1d). The cytotoxicity of the tEF was investigated using an MTT assay on the A2780 human ovarian cancer (OVCA) cell line. The IC_50_ value for dose-dependent growth inhibition of A2780 cells *in vitro* was 128.8 nM when the tEF was well encapsulated and delivered via liposomes, while no significant cytotoxicity of the tEF was observed without transfection (Fig 1a, b), implying that tRNAs or their fragments in the tEF might act as effectors. Furthermore, when rRNA degradation fragments (Fig 1e) were removed from the tEF samples via a Ribo-Zero rRNA removal kit (Illumina), the retained tRNA species induced the death of A2780 cells with increased efficiency (Fig 1f), indicating the major contribution of tRNA species to the overall cytotoxic effects of the tEF. Subsequently, the five most abundant (>500 reads per million) individual tRNAs were purified from small RNA of *T. chinensis* by using a magnetic solid-phase DNA probe method^16^ and were identified by ultra-high performance liquid chromatography coupled with quadrupole time-of-flight mass spectrometry (Fig 1g-i). Cytotoxicity of purified tRNAs was then investigated on A2780 cells, HepG2 of liver cancer and MCF-7 of breast cancer cells. As a result, A2780 and HepG2 cells exhibited a proliferation reduction of over 75% when treated with individual tRNAs at 25 nM (Fig 1j, k), while MCF-7 cells were much less sensitive (reduction of less than 30%) to these tRNA molecules. This pattern corresponded to an approximate IC_50_ value of 14.3 nM for the growth inhibition of A2780 cells caused by treatment with tRNA^Trp(CCA)^ for 48 h, which is close to that of taxol (7.4 nM) used as a positive control drug (Fig 1l). This provides the first evidence of pharmacological activities of plant-derived tRNAs, suggesting that they are a new class of therapeutic agents from medicinal plants.

**Figure 1.**
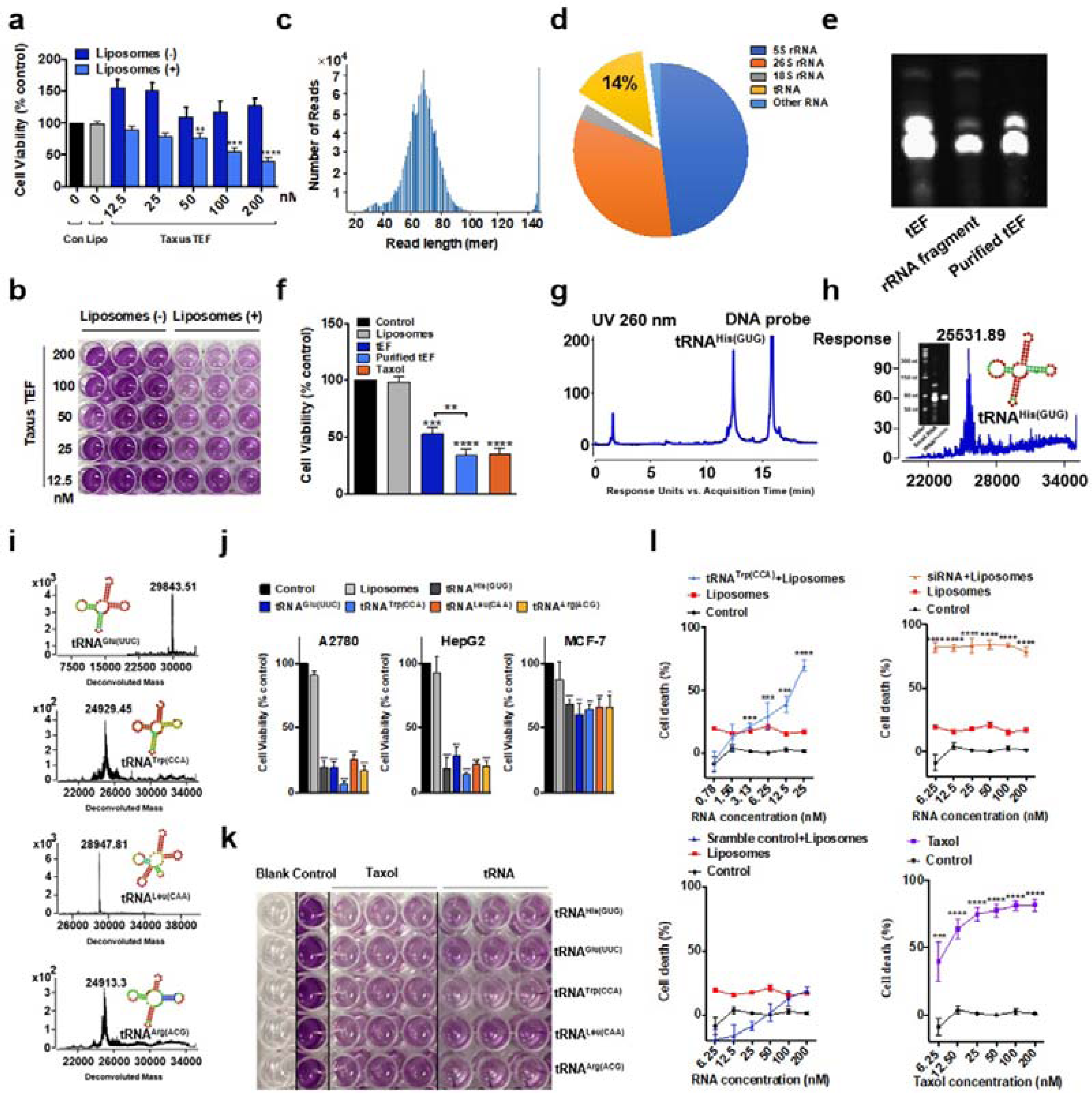
*Taxus*-derived tRNAs exhibit cytotoxicity *in vitro*. **a**, *Taxus* tEF with liposomal transfection exhibited strong cytotoxicity on A2780 cells. **b**, Representative MTT image of the *Taxus* tEF with or without liposomal transfection. **c**, Read length distribution of the tEF determined by NGS was primarily between 60 and 80 mer. **d**, NGS read content analysis showed that 14% of the tEF was tRNA. **e**, Urea-PAGE analysis of the tEF, rRNA fragment and purified tEF. **f**, The cytotoxicity of purified tEF increased after rRNA fragment deletion. **g**, Representative UHPLC method for purification of tRNA^His(GUG)^. **h**, Deconvolution mass spectrometry and urea-PAGE analysis of purified tRNA^His(GUG)^. **i**, Deconvolution mass results of tRNA^Glu(UUC)^, tRNA^Trp(CCA)^. tRNA^Leu(CAA)^ and tRNA^Arg(ACG)^. **j**, MTT assay of the top five abundant tRNAs from Chinese yew on A2780, HepG2 and MCF-7 cells. **k**, Representative MTT image of individual tRNAs on A2780 cells. **l**, Dose-dependent investigation of tRNA^Trp(CCA)^ on A2780 cells. The data are shown as the means ± SDs of 3 independent experiments. \*\**P*<0.01, \*\*\**P*<0.001, \*\*\*\**P*<0.0001.

Meanwhile, since the previous study has revealed that the tRFs could regulate endogenous gene in breast cancer cells^15^, we hypothesized that tRF from the above cytotoxic tRNAs of Chinese yew may be pharmacologically-active. Indeed, tRFs from the 5’ terminal and 3’ ends (5’-tRF and 3’-tRF, respectively) are the most abundant tRFs associated with the AGO protein, which is a key component of the RNA interference pathway^17^. A similar report showed that a 5’-tRF of 19 nucleotides is the most abundant tRF in land plants^18^. Thus, 36 tRFs derived from the top 9 abundant tRNAs of Chinese yew, including 5’-tRF and 3’-tRF with lengths of 22 and 19 nt, respectively, were selected as antisense strands of tRF mimics in siRNA form (Fig 2a, Supplementary Table 2), and the antiproliferative activities of these tRF mimics against A2780 and its taxol-resistant strain (A2780T) cells were screened. The results showed that two of these 36 tRF mimics significantly inhibited the proliferation of the A2780 cells by over 50% at 50 nM (Fig 2b). Among these tRF mimics, tRF-T11 mimic (a double strand RNA with a 22 nt 5’-tRF derived from tRNA^His(GUG)^ as antisense RNA) showed the highest antiproliferative activity against both cell lines. As shown in Fig 2c, the IC_50_ value of tRF-T11 mimic for inhibiting the growth of A2780 cells was 31 nM, which is comparable to that of taxol (10.2 nM). Surprisingly, high sensitivity to tRF-T11 mimic was also observed in A2780T cells, corresponding to an IC_50_ value of 32 nM, which is 80-fold lower than that of taxol (2600 nM). In addition, no significant cytotoxicity of tRF-T11 mimic to IOSE80 normal ovarian surface epithelial cells was observed. Furthermore, the cytotoxicity of tRF-T11 mimic was confirmed by visualizing and tracking fluorescent dye (FAM)-labeled tRF-T11 mimic in A2780 cells. The images shown in Fig 2d indicated that tRF-T11 mimic was efficiently transfected into the cytoplasm of A2780 cells, where they exhibited functions involved in the biological processes, such as suppressing the expression of endogenous targets^19^. These results indicated that *Taxus*-derived tRFs are promising therapeutic agents for drug-resistant OVCA.

**Figure 2.**
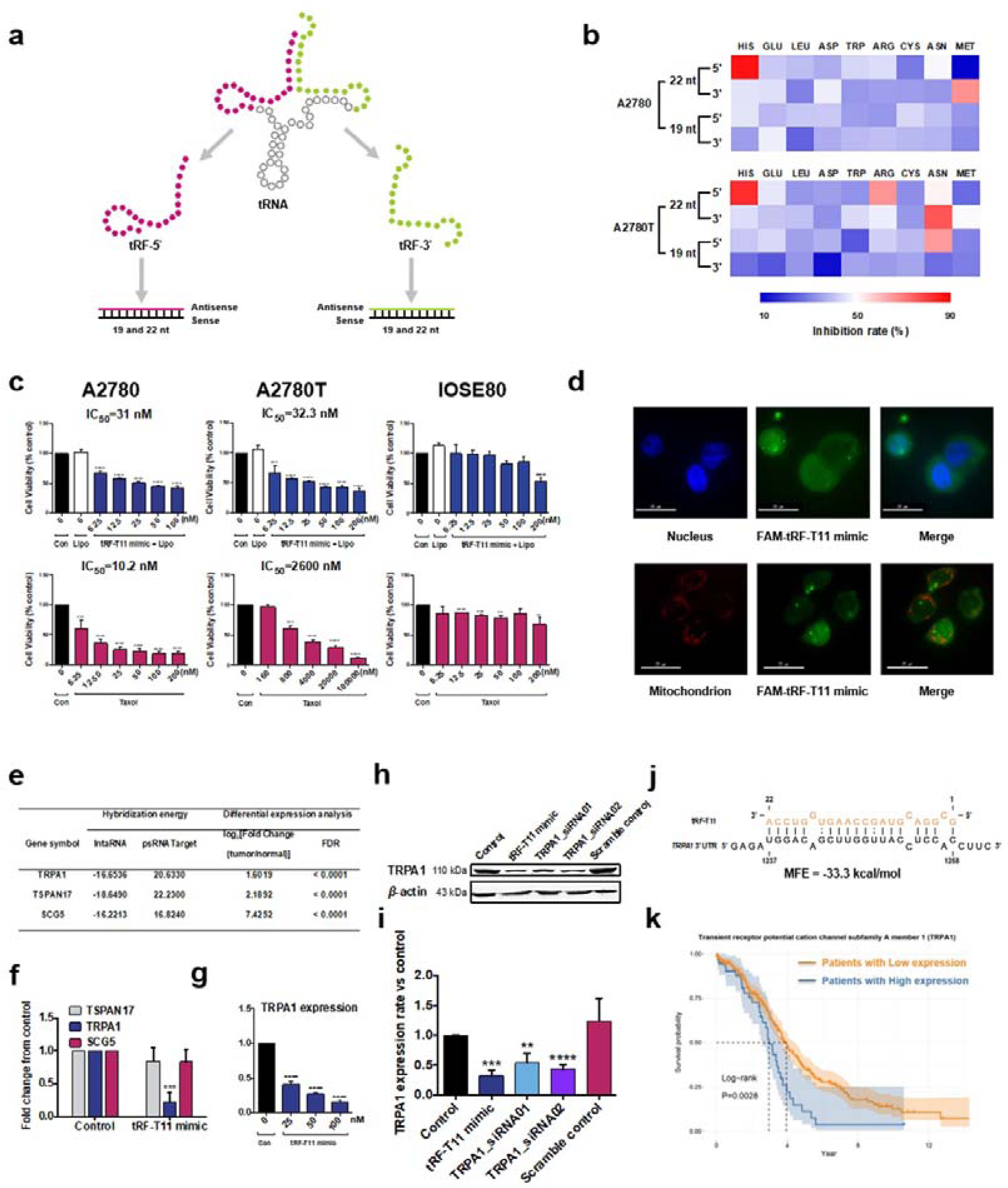
*Taxus*-derived tRF-T11 mimic inhibits human OVCA cell proliferation by targeting *TRPA1 in vitro*. **a**, Design of tRF mimic from 5’ and 3’ end of tRNA. **b**, Heatmap of the inhibition rate of 36 synthesized tRF mimics against A2780 and A2780T cells. **c**, Investigation of the dose dependency of tRF-T11 mimic and taxol against A2780, A2780T and IOSE80 cells. **d**, Fluorescent images capture of FAM labeled tRF-T11 mimic in A2780 cells. **e**, Summary of hybridization energies and differential expression analysis of predicted target genes identified by the IntaRNA and psRNATarget tools. **f**, qPCR validation of the mRNA levels of predicted targets of tRF-T11 in A2780 cells. **g**, tRF-T11 mimic decreased the mRNA levels of *TRPA1* in a dose-dependent manner. **h**, Western blot assay for the evaluation of *TRPA1* expression in A2780 cells treated with tRF-T11 mimic, siRNA against *TRPA1* (TRPA1_siRNA01, TRPA1_siRNA02) and scrambled control at 25 nM. Band intensities were measured using ImageJ and were normalized to those of β-actin (**i**). **j**, Putative tRF-T11 binding sites in the *TRPA1* 3’ UTR. **k**, Overall survival probability of patients in the TCGA OVCA cohort. The data are shown as the means ± SDs of 3 independent experiments. \*\**P*<0.01, \*\*\**P*<0.001, \*\*\*\**P*<0.0001.

The divergent activity of tRF mimics derived from individual tRNAs observed in this study suggested that plant-derived tRFs can exhibit bioactivities in a sequence-specific manner in mammalian cells. It has been reported that functionality of exogenous tRFs possibly overlaps that of miRNAs, a characteristic that has been widely documented for endogenous tRFs in both plants and mammalian cells^20, 21^. To further identify potential target genes of tRF-T11, we performed an *in silico* prediction using two independent bioinformatic tools, IntaRNAv2^22^ and psRNATarget^23^, followed by differential expression analysis based on the gene expression profiles in normal and tumor ovarian tissue to prioritize the candidate targets. As a result, the top three differentially expressed genes, namely transient receptor potential cation channel subfamily A member 1 (*TRPA1*), tetraspanin 17 (*TSPAN17*) and secretogranin V (*SCG5*), with the minimum required energy to hybridize with tRF-T11 at their 3’ UTR regions of mRNA, were considered as the potential targets for further experimental validation (Fig 2e). Among these three genes, *TRPA1*, a new target recently unveiled by Takahashi *et al*^24^ for cancer therapies, which acts by triggering a noncanonical oxidative stress defense mechanism of tumor suppression, was dramatically downregulated in A2780 cells (by approximately 75%) in a dose-dependent manner by tRF-T11 mimic treatment (Fig 2j, g), while the mRNA levels of the other two targets did not vary significantly (Fig 2f). Furthermore, a marked decrease of over 50% in the protein expression level of *TRPA1* was also observed in A2780 cells treated with tRF-T11 mimic (Fig 2h, i), which may act as a translational repressor of the *TRPA1* gene by acting like a miRNA in human OVCA cells. Indeed, previous reports revealed that *TRPA1* overexpression is associated with poor survival probability in patients (Fig 2k). Unfortunately, no clinical therapeutic drug targeting *TRPA1* is available, and our findings provide a novel RNA sequence derived from plant can potently silence human *TRPA1*.

Recently, advances in RNA therapeutics have been facilitated by breakthroughs in pharmaceutical delivery systems, including liposomes, nanoparticles, dendrimers and carbon nanotubes have been shown to provide promising application *in vitro* and *in vivo*^25^. To evaluate the antitumor activity *in vivo*, a histidine-lysine polymer (HKP) nanoparticle system was applied to deliver tRF-T11 mimic to a well-established xenograft mouse model of human OVCA cells. The mass ratio of HKP and tRF-T11 mimic was optimized (Fig 3a, b) with an appropriate particle size (164.2 nm) and good dispersion (41.2 mV, Fig 3c). As shown in Fig 3d to g, compared to vehicle control group, tRF-T11 mimic treatment of xenografts for 20 days significantly diminished the tumor growth rate by approximately 50% at a dose of only 177 nmol/kg, which is more than 16-fold lower than that of taxol (2928 nmol/kg) required to achieve comparable tumor-suppressive effects without significant adverse impacts on body weight. Compared to those from control mice, the protein expression levels of *TRPA1* in the tumors with intratumoral injection of high-dose tRF-T11 mimic exhibited significant reduction (Fig 3h, i), which is consistent with the western blot results *in vitro*. Although taxol is a commonly used first-line drug for cancer patients, chemoresistance is usually observed. Since tRF-T11 mimic exhibited comparable tumor suppression efficacy to taxol at a much lower dose, our study may indicate an alternative treatment option.

**Figure 3.**
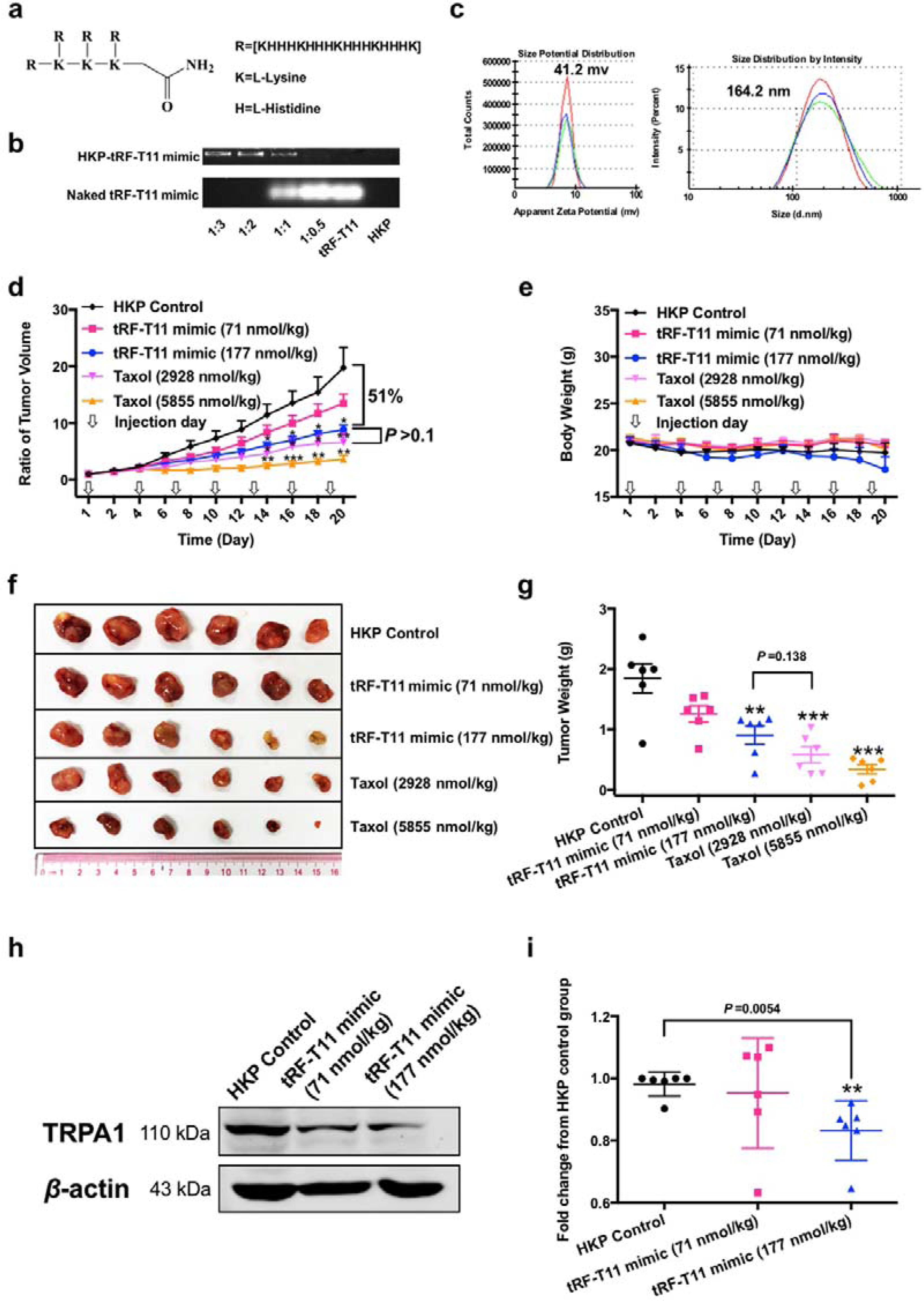
*Taxus*-derived tRF-T11 mimic decreases OVCA tumor growth via targeting *TRPA1 in vivo*. **a**, Structure of HKP. **b**, Optimization of the mass ratio of tRF-T11 mimic and HKP. **c**, The particle sizes and zeta potential of the HKP-tRF-T11 mimic. **d**, Growth rate of tumors treated with tRF-T11 mimic, taxol and HKP alone. **e**, Quantification of body weights in each group. **f**, Images of each tumor. **g**, No significant difference in tumor weight was observed during treatment in the high-dose tRF-T11 mimic group and the low-dose taxol group. **h**, Representative western blot image of *TRPA1* protein levels in tumors of different treatment. **i**, Band intensity ratio of *TRPA1*/β-actin in tumors treated with different doses of tRF-T11 mimic (n=6 per group). The data are shown as the means ± SDs of 3 independent experiments excluding those shown in **c** (± SEM). \**P*<0.05, \*\**P*<0.01, \*\*\**P*<0.001.

In summary, this study provides the first evidence that plant-derived tRNAs and tRFs from Chinese yew exhibit potent antitumor effects against human OVCA, which will greatly expand our knowledge on the active ingredients in natural sources, and provide additional clues for evaluating the active components of medicinal plants or herbal medicines. Although cross-kingdom regulation in human by miRNA uptake from food and medicinal plants has been widely studied, this report emphasizes the potential regulatory roles of plant-derived small RNAs from an abundance of housekeeping noncoding RNA species, such as tRNAs and tRFs. This infinitely vast pool filled with RNA sequences and chemical modifications provided by nature can serve as a new sequence library for the development of novel RNAi-based therapeutics. Furthermore, since chemical modified oligonucleotides could be simply and efficiently synthesized in large amount using the phosphoramidite methodology^26, 27^, this discovery demonstrated a pilot strategy for identifying pharmacologically active tRFs from plants, such as those from the genus *Taxus*.

As a promising and rapidly advancing frontier, siRNA can be readily designed based on a well-validated target mRNA. Often, the process of unraveling the pathogenesis of many human diseases to find key molecular targets for drug treatment is time-consuming and difficult without a functional approach with bottom-up guidance. As a result, the effective design of RNAi therapeutic drug candidates will be difficult or impossible due to the lack of reliable targets. As a benefit of the high target specificity and selectivity of RNA drugs, pharmacologically active RNA molecules identified from medicinal plants could, in turn, lead to the improvement of the efficiency and possibilities of discovering new drug target, and shed light on disease etiology.

## Acknowledgements

We thank Novogene Bioinformatics Institute (Beijing, China) for technical support in next-generation sequencing; Y. L. (Sirnaomics, Inc., U.S.A.) for technical advice and the gift of HKP for RNA delivery *in vivo*. This work was financially funded by the CUHK Direct Grant 4053364, the Innovation and Technology Commission, Hong Kong Government to the State Key Laboratory, and grant from The Science and Technology Development Fund, Macau SAR (File no. 015/2017/AFJ).

## Contributions

Z.-H.J. and T.-M.Y. conceived the study. K.-Y.C., T.-M.Y. and T.-F.C. designed the experiments. K.-Y.C., T.-M.Y., J.-Z.Z. and J. L. conducted the experiments. K.-Y.C., J.-Z.Z. and C. L. analyzed the data. K.-Y.C., T.-M.Y. and E.L.L. drafted the manuscript. All authors approved the manuscript.

## Methods

### Plant materials and RNA extraction of *Taxus chinensis*

Plants of *Taxus chinensis (Pilger) Rehd. var. mairei* (Lemee et Levl.) Cheng et L. K. Fu used in this study were collected from Sanming City, Fujian Province, China. Plant materials were freshly cut, frozen and stored in liquid nitrogen until use. Plant tissues were ground into a fine powder in liquid nitrogen and were then homogenized (IKA, Germany) in CTAB buffer. After incubation for 2 min at 65°C, the tissue lysate was cooled immediately on ice for 10 min, followed by centrifugation at 12,000 ×g for 15 min at 4°C. The supernatant was collected and extracted with an equal volume of phenol:chloroform:isopentanol (50:48:1). Phases were separated at 4°C by centrifugation at 12,000 ×g for 15 min. The supernatant was extracted again as described above with chloroform:isopentanol (24:1) and mixed with an equal volume of 6 M guanidinium thiocyanate solution (Tokyo Chemical Industry, Japan) and ethanol (Anaqua Chemicals Supply, U.S.A.) to a final concentration of 55%. The mixture was passed through a silicon-membrane column and washed with 80% ethanol. Finally, total RNA was eluted with RNase-free water, and small RNA species were enriched by using a mirVana miRNA isolation kit (Invitrogen, U.S.A.) following the manufacturer’s instructions.

### tEF preparation

Small RNA was separated by 6% urea-polyacrylamide gel electrophoresis (PAGE). Gel fractions containing the tEF, as visualized by SYBR Gold nucleic acid gel staining (Thermo, U.S.A.), were excised and electroeluted in 3 kD molecular weight cut off dialysis tubing (Spectrum, U.S.A.) at 100 V for 1.5 h. The tEF was finally desalted and recovered on a silicon-membrane column.

### tEF library construction and sequencing

Sequencing libraries were generated by using a TruSeq small RNA Library Preparation Kit (Illumina, U.S.A.), followed by a round of adaptor ligation, reverse transcription and PCR. PCR products were then purified, and libraries were quantified on an Agilent Bioanalyzer 2100 system (Agilent Technologies, U.S.A.). The library preparations were sequenced at the Novogene Bioinformatics Institute (Beijing, China) on an Illumina HiSeq platform using the 150-bp paired-end (PE150) strategy to generate over 15 million raw paired reads. A total of 1,729,438 clean reads were obtained by removing low quality regions and adaptor sequences. The tRNA genes were identified by using the tRNAscan-SE 2.0 program (http://lowelab.ucsc.edu/tRNAscan-SE/) and annotated by searching the Nucleotide Collection (nr/nt) database using the Basic Local Alignment Search Tool (BLAST) program (https://blast.ncbi.nlm.nih.gov/Blast.cgi).

### Purification of individual tRNAs

Synthetic single-stranded DNA oligonucleotides (BGI, China; Supplementary Table 3) complementary to the target tRNA were mixed with *Taxus* small RNA and denatured at 95°C for 5 min, followed by incubation at a temperature 5°C lower than the melting temperature (Tm) for 1.5 h. Streptavidin magnetic beads (Beaverbio, China) were then added to the mixture and incubated for 30 min at the annealing temperatures. Then, the biotinylated DNA/tRNA-coated beads were separated with a magnet and washed at 40°C. Subsequently, the magnetic beads were resuspended in RNase-free water, and the immobilized tRNA with probes was released by incubation at 65°C for 5 min. The solutions were directly injected using an ACQUITY UPLC OST C_18_ column (2.1×100 mm i.d., 1.7 μm; Waters, U.S.A.) maintained at 60°C into a UHPLC Agilent 1290 system (Agilent, U.S.A.) equipped with a diode array detector. The flow rate was set at 0.2 mL/min, and a gradient elution was performed with (A) 100 mM HFIP+15 mM TEAA and (B) 50% MeOH in A as follows: 0-1.5 min, 2% B; 1.5-8.3 min, 2%-28% B; and 8.3-16.5 min, 28%-34% B. MS data were acquired on an Agilent 6545 Q-TOF mass spectrometer with a gas temperature of 320°C, spray voltage of 3.5 kV, and sheath gas flow and temperature of 12 L/min and 350°C, respectively. For the MS experiment, samples were analyzed in negative polarity over an m/z range of 500 to 3200. Fractions from chromatographic peaks of individual tRNAs were collected and freeze dried.

### RNA transfection and cytotoxicity determination

RNA samples dissolved in RNase-free water were stored at −20°C before use. Transfection of RNA samples was performed according to the manufacturer’s instructions for Lipofectamine RNAiMAX Transfection Reagents (Thermo, U.S.A.). After treatment for 48 h, an MTT assay was performed, and absorbance values were calorimetrically determined at 570 nm using a SpectraMax 190 microplate reader (Molecular Devices, Sunnyvale, U.S.A). The dose-dependent IC_50_ values were calculated by GraphPad Prism 5.0 (GraphPad, U.S.A.). A siRNA (Forward: 5’-GGAGCGAUUUAGCCAAGAATT-3’, Sense: 5’-UUCUUGGC UAAAUCGCUCCTT-3’) was used as positive control. Each experiment was carried out three times. The results are expressed as the means ± standard deviations.

### Acquisition of fluorescence images of transfected FAM-labeled tRF-T11 mimic in A2780 cells

Briefly, 1×10^5^ A2780 cells were seeded in glass bottom cell culture dishes (Nest Biotechnology, China). The medium was discarded after 20 h, and 50 nM FAM-labeled tRF-T11 mimic (GenePharma, China) was transfected into the cells. After 6 h, the transfection medium was discarded, and the cells were rinsed 3 times with RPMI 1640 medium. For nuclear staining, cells were incubated with 1 mL of Hoechst 33258 (Sigma, U.S.A.) for 10 min at 37°C, followed by washing with RPMI 1640 medium. For mitochondrial staining, cells were incubated with 1 mL of MitoBeacon Red (GeneCopoeia, U.S.A.) for 30 min at 37°C, followed by washing with RPMI 1640 medium. Fluorescence images were acquired with an API DeltaVision Live-cell Imaging System (Applied Precision Inc., GE Healthcare Co., U.S.A.) and were captured at x60 magnification, scale bar, 25 µm.

### tRF-T11 target prediction and prioritization

Previously described small RNA or tRF target prediction methods and restriction criteria were applied in this study to identify functional target genes of tRF-T11^28^. Specifically, the 3’ UTR sequences of all mRNAs retrieved from GENCODE (v23) and the sequence of tRF-T11 derived from *Taxus* were firstly loaded into IntaRNAv2 and psRNATarget to predict the potential targets. The top 10 overlapping targets predicted were then prioritized and filtered by crosschecking their expression in tumor and normal ovarian tissues. Accordingly, we first obtained the gene expression profiles for 379 OVCA tissues and 88 normal ovarian tissues from UCSC Xena, which contains curated comprehensive batch-effect-adjusted pancancer genomic data^29^. Subsequently, EBSeq^30^ was employed to filter the target genes differentially expressed between tumor and normal ovarian tissues. The predicted targets satisfying two criteria, fold change (tumor/normal) > 2 and FDR < 0.1.

### Quantitative real-time PCR

Total RNA was extracted from A2780 cells treatment with the tRF-T11 mimic for 48 h using TRIzol (Invitrogen, U.S.A.) on a silicon-membrane column. RNA was then reverse transcribed using the GoScript Reverse Transcription System (Promega, U.S.A.). Quantitative real-time PCR was performed on a high-productivity real-time quantitative PCR ViiA^TM^ 7 system (Life Technologies, U.S.A.) using GoTaq^®^ qPCR Master Mix (Promega, U.S.A.) according to the manufacturers’ instructions. The PCR primers were synthesized commercially (BGI, China, Supplementary Table 4). All data are presented as the mean fold change values duplicated and normalized to β-actin expression using the 2^−CT^ method. Each experiment was carried out three times. The results are expressed as the means ± standard deviations.

### Western blot analysis

A2780 cell lysates were collected after treatment with tRF-T11 mimic for 48 h, and protein extraction was carried out using RIPA buffer (Cell Signaling Technology, U.S.A.) supplemented with protease and phosphatase inhibitor cocktails (Roche, Switzerland). Protein concentrations were then determined by a BCA protein assay (Thermo, U.S.A.). Denatured proteins were separated by 8% SDS-PAGE and transferred to nitrocellulose membranes, followed by blocking with 5% bovine serum albumin (Thermo, U.S.A.) for 2 h. Membranes containing proteins were incubated with primary antibody against *TRPA1* (1:500 dilution, Novus, U.S.A.) or β-actin (1:1000 dilution, Santa Cruz, U.S.A.) overnight, followed by washing in TBST and incubation with a fluorescent secondary antibody (Li-cor, U.S.A.) for detection. Band intensities were measured using ImageJ and were normalized to those of β-actin. The sequences of TRPA1_siRNA01, TRPA1_siRNA02 and the scrambled control (GenePharma) are shown in Supplementary Table 5. Each experiment was carried out three times. The results are expressed as the means ± standard deviations.

### Preparation of HKP-tRF-T11 mimic nanoparticles

HKP (a gift from Dr. Yang Lu, Sirnaomics, Inc., U.S.A.) dissolved in RNase-free water (240 μg/mL) were first balanced overnight at 4°C. Equal volumes of tRF-T11 mimic solutions at a concentration of 80 μg/mL were then mixed and incubated at room temperature for 30 min. The particle sizes and zeta potential of the resulting nanoparticles were then evaluated by a Malvern Zetasizer Nano ZS90 (Malvern Instruments, U.K.).

### *In vivo* models for tumor therapy

A2780 cells (4.0×10^6^) were injected subcutaneously into the armpits of 8-week-old BALB/c nude mice (Shanghai SLAC Laboratory Animal Co., Ltd., China). When the tumors reached 50 mm^3^, the tumor-bearing nude mice were treated with different doses (71 nmol/kg, 177 nmol/kg) of tRF-T11 mimic (GeneDesign, Inc. Osaka, Japan) encapsulated with HKP and different doses of taxol (2928 nmol/kg or 5855 nmol/kg) via intratumoral injection once every three days. A group treated with HKP alone was established as a control. The animals were sacrificed on day 20. Tumor diameters were measured at the points of maximum length and maximum width with digital calipers. Tumor volumes were calculated by the following formula: volume=(width)^2^×length/2. The data were statistically analyzed using GraphPad Prism 5.0. Animal care and use protocols were performed in compliance with the guidelines of the Ethics Committee of the College of Pharmaceutical Sciences, Southwest University.

