## Supplementary Figure for "Cross-kingdom regulation of tRNAs/tRFs derived from Chinese yew"

**Supplementary Figure Legends**

**Supplementary Figure 1.** **Total RNA in good quality were extracted from *Taxus chinensis***. **a**, Nanodrop indicated that extracted total RNA is in high purity. **b,** Scalable total RNA extraction using a developed CTAB-based method. **c**, Separation and enrichment of small RNA from large RNA.

**Supplementary Figure 2.** **Quality evaluation confirmed that small RNA was well separated from large RNA**. **a**. Agilent 2100 Bioanalyzer profile of small RNA and large RNA from *Taxus chinensis*. **b**. Monitored electrophoresis gram of small RNA and large RNA.

**Supplementary Figure 3.** **Preparation of** **tEF from small RNA of Chinese yew**. **a**. Small RNA was gel-fractionated and tEF was sliced off followed by electroelution. **b**. Agilent 2100 Bioanalyzer showed that compared to small RNA, 5S, 5.8S rRNA and other RNA species could not be detected in tEF.

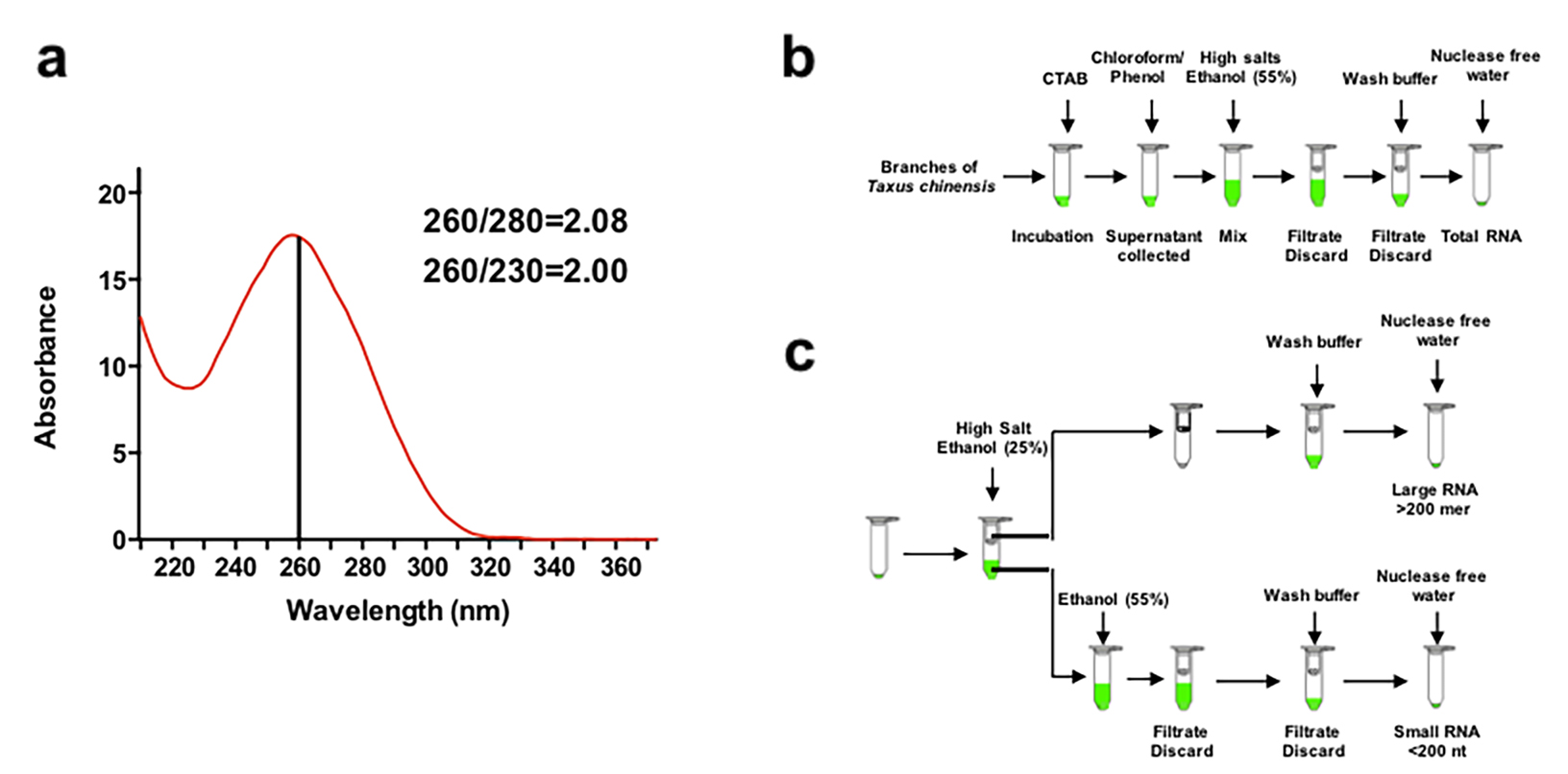

**Supplementary Figure 1.**

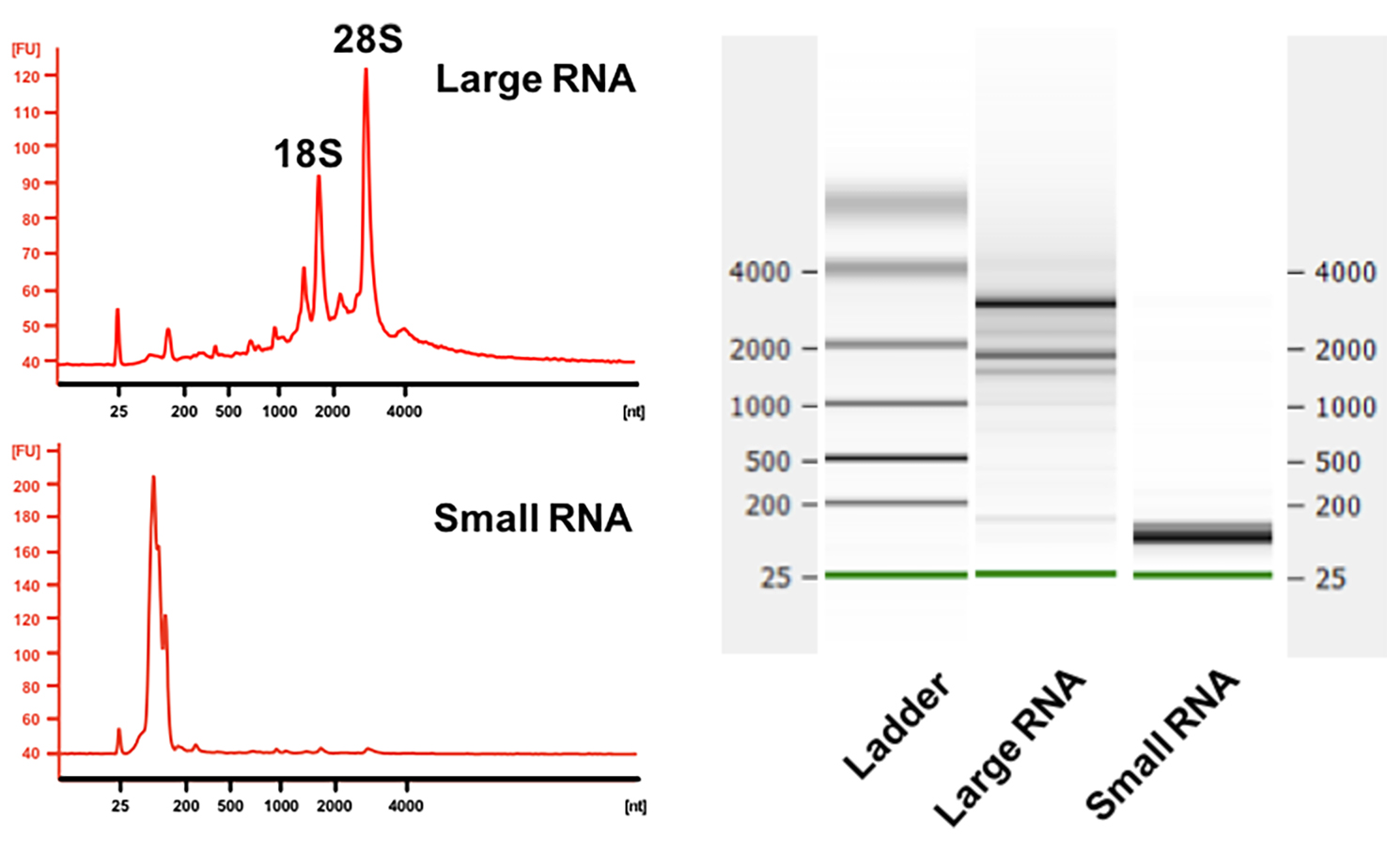

**Supplementary Figure 2.**

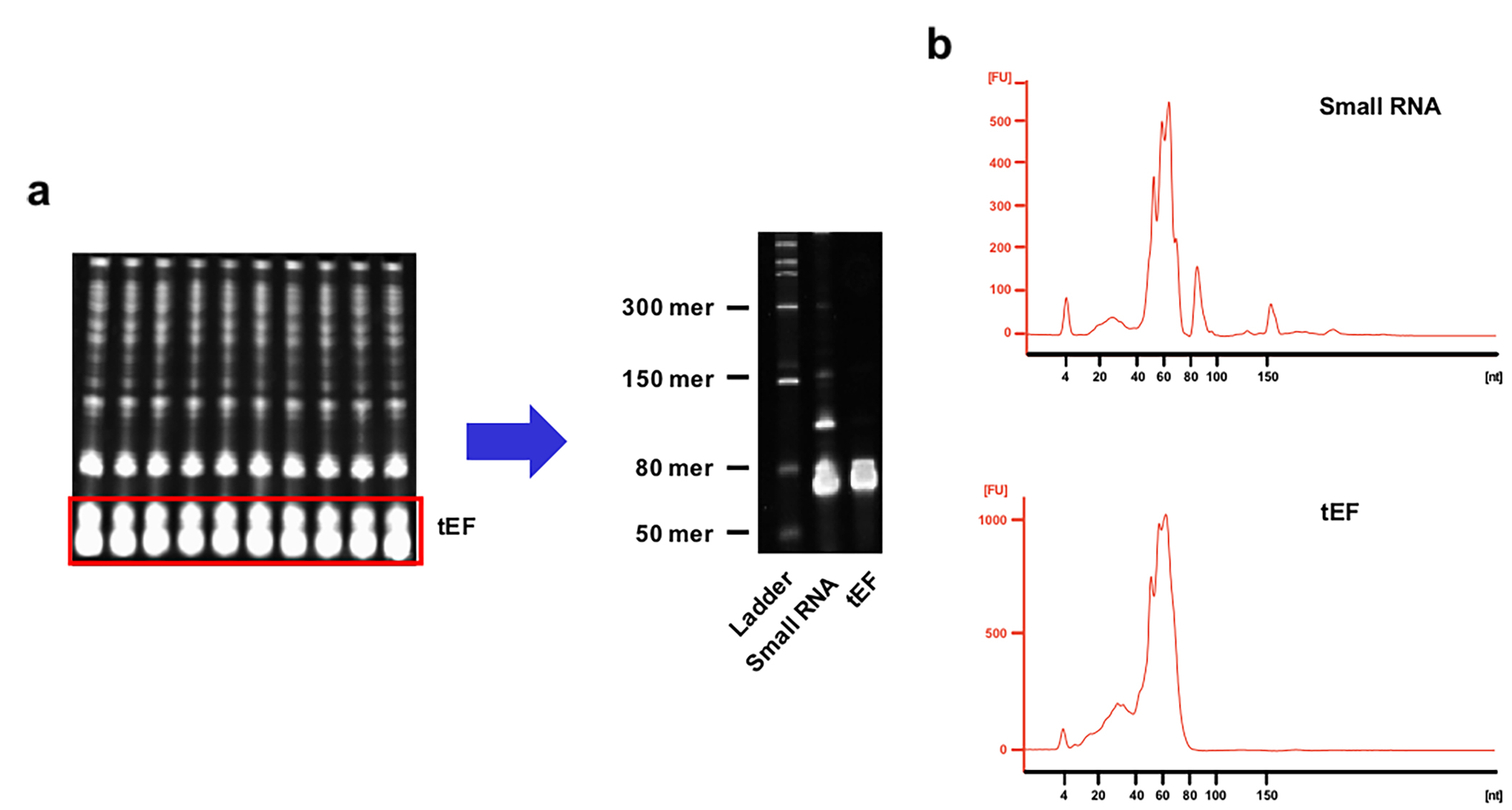

**Supplementary Figure 3.**
