## Supplementary Table 1 for "Cross-kingdom regulation of tRNAs/tRFs derived from Chinese yew"

**Supplementary Table 1. Identified tRNA sequences in Chinese yew by NGS**

| tRNA | Sequence (5'-3') | Length (mer) |
| --- | --- | --- |
| tRNA^His(GUG)^ | GCGGACGUAGCCAAGUGGUCCAAAGGCAGUGGAUUGUGAAUCCACCACGCGCGGGUUCAAUCCCCGUCGUUCGCCCCA | 78 |
| tRNA^Glu(UUC)^ | GCCCCUAUCGUCUAGUGGCCCAGGACAUCUCUCUUUCAAGGAGGCAACGGGGAUUCGAUUUCCCCUAGGGGUACCA | 76 |
| tRNA^Trp(CCA)^ | GCGCUCUUAGUUCAGUGCGGUAGAACGCAGGUCUCCAAAACCUGAUGCCGUAGGUUCAAAUCCUACAGAGCGCCA | 75 |
| tRNA^Leu(CAA)^ | GCCUUGAUGGUGAAAUGGUAGACACGCGAGACUCAAAAUCUCGUGCUAAACAGCGUGGAGGUUCGAAUCCUCUUCAAGGCACCA | 84 |
| tRNA^Arg(ACG)^ | GGGCCUGUAGCUCAGAGGAUUAGAGCACGUGGUUGCGAACCACGGUGUCGGGGGUUCGAAUCCCUCCUCGCCCACCA | 77 |
| tRNA^Asp(GUC)^ | GGGAUUGUAGUUCAAUUGGUUAGAGUACCGCCCUGUCAAGACGGAAGUUGCGGGUUCGAGCCCCGUCAGUCCCGCCA | 77 |
| tRNA^Asn(GUU)^ | UCCUCAGUAGCUCAGUGGUAGAGCGGUCGGCUGUUAACCGAUUGGUCGUAGGUUCAAAUCCUAUUUGAGGAGCCA | 75 |
| tRNA^Cys(GCA)^ | GGCGACAUAGCCAAGUGGUAAGGCAGGGGACUGCAAAUCCCCCAUCCCCAGUUCAAAUCCGGGUGUCGCCUCCA | 74 |
| tRNA^Gln(UUG)^ | GGGGCGUGGCCAAGCGGUAAGGCAACAGGUUUUGGUCCUGUUAUUGCGAAGGUUCGAAUCCUUUCGUCCCAGCCA | 75 |
| tRNA^Gly(GCC)^ | GGGUAUUGUUUAAUGGAUAAAAUUUAUUCUUGCCAAGGAUAAGAUGCGGGUUCGAUUCCCGCUACCCGCCCA | 72 |
| tRNA^IIe(UAU)^ | AGGGAUAUAACUCAGUAGUAGAGUGUCACCUUUAUGUGGUGAAAGUCAUCAGUUCAAACCUGAUUAUCCCUACCA | 75 |
| tRNA^Leu(UAG)^ | GCCGCCAUGGUGAAAUUGGUAGACACGCUGCUCUUAGGAAGCAGUGCUAGAGCAUCUCGGUUCGAAUCCGAGUGGUGGCACCA | 83 |
| tRNA^Leu(UAA)^ | GGGGAUAUGGCGGAAUUGGUAGACGCUACGGACUUAAAAAAUCCGUUGGUUUUAUAAACCGUGAGGGUUCAAGUCCCUCUAUCCCCACCA | 90 |
| tRNA^Lys(UUU)^ | GGGUUGUUAACUCAAUGGUAGAGUACUCGGCUUUUAACCGAcGAGUUCCGGGUUCAAGUCCCGGGCAACCCACCA | 75 |
| tRNA^Met(CAU)^ | GCAUCCAUGGCUGAAUGGUCAAAGCACCCAACUCAUAAUUGGGAAGUCGCGGGUUCAAUUCCUGCUGGAUGCACCA | 76 |
| tRNA^Met(CAU)^ | CGCGGAGUAGAGCAGUUUGGUAGCUCGCAAGGCUCAUAACCUUGAAGUCACGGGUUCAAAUCCCGUCUCCGCAACCA | 77 |
| tRNA^Phe(GAA)^ | GUCGGGAUAGCUCAGUUGGUAGAGCAGAGGACUGAAAAUCCUCGUGUCACCAGUUCAAAUCUGGUUCCUGGCACCA | 76 |
| tRNA^Pro(UGG)^ | AGGGAUGUAGCGCAGCUUGGUAGCGCGUUUGUUUUGGGUACAAAAUGUCGCAGGUUCAAAUCCUGUCAUCCCUACCA | 77 |
| tRNA^Pro(GGG)^ | CGGAGCAUAACGCAGUUUGGUAGCGUGCCAUCUUGGGGUGAUGGAGGUCGCGGGUUCAAAUCCUGUUGCUCCGACCA | 77 |
| tRNA^Ser(UGA)^ | GGAGAGAUGGCCGAGUGGUUGAUGGCUCCGGUCUUGAAAACCGGUAUAGUUUUAAAAACUAUCGAGGGUUCGAAUCCCUCUCUCUCCUCCA | 91 |
| tRNA^Ser(GCU)^ | GGAGAGAUGGCUGAGCGGACUAAAGCGGUGGAUUGCUAAUCCGUUGUACAGACUAUCUGUACCGAGGGUUCGAAUCCCUCUUUCUCCGCCA | 91 |
| tRNA^Thr(UGU)^ | GCCUGCUUAGCUCAGAGGUUAGAGCAUCGCACUUGUAAUGCGACGGUCAUCGGUUCGAUCCCGAUAGAAGGCUCCA | 76 |
| tRNA^Thr(GGU)^ | GCACUUUUAACUCAGUGGUAGAGUAACGCCAUGGUAAGGCGUAAGUCAUCGGUUCAAGCCCGAUAAAGGGCUCCA | 75 |
| tRNA^Tyr(GUA)^ | GGGUCGAUGCCCGAGUGGCUAAUGGGGACGGACUGUAAAUCCGUUGGCAAUAUGCUUACGCUGGUUCAAAUCCAGCUCGGCCCACCA | 87 |
| tRNA^Arg(CUC)^ | GCGUCCAUCGUCUAAUGGAUAGGACAGAGGUCUUCUAAACCUUAGGUAUAGGUUCAAAUCCUAUUGGACGUACCA | 75 |
