## Supplementary Table 2 for "Cross-kingdom regulation of tRNAs/tRFs derived from Chinese yew"

**Supplementary Table 2. Synthetic tRF mimics from top 9 abundant tRNAs in Chinese yew**

| Source | Code | Antisense derived from tRNA (5'-3') | Sense (5'-3') | Length (mer) | Class |
| --- | --- | --- | --- | --- | --- |
| tRNA^His(GUG)^ | tRF-T11 | GCGGACGUAGCCAAGUGGUCCA | UGGACCACUUGGCUACGUCCGC | 22 | 5' |
|  | tRF-T20 | GCGGACGUAGCCAAGUGGU | ACCACUUGGCUACGUCCGC | 19 |  |
|  | tRF-T12 | UCAAUCCCCGUCGUUCGCCCCA | UGGGGCGAACGACGGGGAUUGA | 22 | 3' |
|  | tRF-T42 | AUCCCCGUCGUUCGCCCCA | UGGGGCGAACGACGGGGAU | 19 |  |
| tRNA^Glu(UUC)^ | tRF-T16 | GCCCCUAUCGUCUAGUGGCCCA | UGGGCCACUAGACGAUAGGGGC | 22 | 5' |
|  | tRF-T25 | GCCCCUAUCGUCUAGUGGC | GCCACUAGACGAUAGGGGC | 19 |  |
|  | tRF-T17 | UCGAUUUCCCCUAGGGGUACCA | UGGUACCCCUAGGGGAAAUCGA | 22 | 3' |
|  | tRF-T43 | AUUUCCCCUAGGGGUACCA | UGGUACCCCUAGGGGAAAU | 19 |  |
| tRNA^Trp(CCA)^ | tRF-T30 | GCGCUCUUAGUUCAGUGCGGUA | UACCGCACUGAACUAAGAGCGC | 22 | 5' |
|  | tRF-T23 | GCGCUCUUAGUUCAGUGCG | CGCACUGAACUAAGAGCGC | 19 |  |
|  | tRF-T31 | GUUCAAAUCCUACAGAGCGCCA | UGGCGCUCUGUAGGAUUUGAAC | 22 | 3' |
|  | tRF-T46 | CAAAUCCUACAGAGCGCCA | UGGCGCUCUGUAGGAUUUG | 19 |  |
| tRNA^Leu(CAA)^ | tRF-T18 | GCCUUGAUGGUGAAAUGGUAGA | UCUACCAUUUCACCAUCAAGGC | 22 | 5' |
|  | tRF-T22 | GCCUUGAUGGUGAAAUGGU | ACCAUUUCACCAUCAAGGC | 19 |  |
|  | tRF-T19 | UCGAAUCCUCUUCAAGGCACCA | UGGUGCCUUGAAGAGGAUUCGA | 22 | 3' |
|  | tRF-T44 | AAUCCUCUUCAAGGCACCA | UGGUGCCUUGAAGAGGAUU | 19 |  |
| tRNA^Arg(ACG)^ | tRF-T32 | GGGCCUGUAGCUCAGAGGAUUA | UAAUCCUCUGAGCUACAGGCCC | 22 | 5' |
|  | tRF-T24 | GGGCCUGUAGCUCAGAGGA | UCCUCUGAGCUACAGGCCC | 19 |  |
|  | tRF-T33 | UCGAAUCCCUCCUCGCCCACCA | UGGUGGGCGAGGAGGGAUUCGA | 22 | 3' |
|  | tRF-T47 | AAUCCCUCCUCGCCCACCA | UGGUGGGCGAGGAGGGAUU | 19 |  |
| tRNA^Asp(GUC)^ | tRF-T28 | GGGAUUGUAGUUCAAUUGGUUA | UAACCAAUUGAACUACAAUCCC | 22 | 5' |
|  | tRF-T21 | GGGAUUGUAGUUCAAUUGG | CCAAUUGAACUACAAUCCC | 19 |  |
|  | tRF-T29 | UCGAGCCCCGUCAGUCCCGCCA | UGGCGGGACUGACGGGGCUCGA | 22 | 3' |
|  | tRF-T45 | AGCCCCGUCAGUCCCGCCA | UGGCGGGACUGACGGGGCU | 19 |  |
| tRNA^Cys(GCA)^ | tRF-T34 | GGCGACAUAGCCAAGUGGUAAG | CUUACCACUUGGCUAUGUCGCC | 22 | 5' |
|  | tRF-T26 | GGCGACAUAGCCAAGUGGU | ACCACUUGGCUAUGUCGCC | 19 |  |
|  | tRF-T35 | UCAAAUCCGGGUGUCGCCUCCA | UGGAGGCGACACCCGGAUUUGA | 22 | 3' |
|  | tRF-T48 | AAUCCGGGUGUCGCCUCCA | UGGAGGCGACACCCGGAUU | 19 |  |
| tRNA^Asn(GUU)^ | tRF-T36 | CCUCAGUAGCUCAGUGGUAGAG | CUCUACCACUGAGCUACUGAGG | 22 | 5' |
|  | tRF-T27 | CCUCAGUAGCUCAGUGGUA | UACCACUGAGCUACUGAGG | 19 |  |
|  | tRF-T37 | GGUUCAAAUCCUAUUUGAGGAG | CUCCUCAAAUAGGAUUUGAACC | 22 | 3' |
|  | tRF-T49 | UCAAAUCCUAUUUGAGGAG | CUCCUCAAAUAGGAUUUGA | 19 |  |
| tRNA^Met(CAU)^ | tRF-T38 | CGCGGAGUAGAGCAGUUUGGUA | UACCAAACUGCUCUACUCCGCG | 22 | 5' |
|  | tRF-T40 | CGCGGAGUAGAGCAGUUUG | CAAACUGCUCUACUCCGCG | 19 |  |
|  | tRF-T39 | GGUUCAAAUCCCGUCUCCGCAA | UUGCGGAGACGGGAUUUGAACC | 22 | 3' |
|  | tRF-T41 | UCAAAUCCCGUCUCCGCAA | UUGCGGAGACGGGAUUUGA | 19 |  |
