## Supplementary Table 3 for "Cross-kingdom regulation of tRNAs/tRFs derived from Chinese yew"

**Supplementary Table 3. DNA probes for purification of individual tRNAs**

| tRNA | Probe (5'-3') |
| --- | --- |
| tRNA^His(GUG)^ | GGCGAACGACGGGGATTGAACCCGCGCGTG |
| tRNA^Glu(UUC)^ | TTGCCTCCTTGAAAGAGAGATGTCCTGGGC |
| tRNA^Trp(CCA)^ | ACGGCATCAGGTTTTGGAGACCTGCGTTCT |
| tRNA^Leu(CAA)^ | ACGCTGTTTAGCACGAGATTTTGAGTCTCG |
| tRNA^Arg(ACG)^ | CGTGGTTCGCAACCACGTGCTCTAATCCTC |
