## Supplementary Table 4 for "Cross-kingdom regulation of tRNAs/tRFs derived from Chinese yew"

**Supplementary Table 4. Primers for quantitative real-time PCR**

| Gene name | Forward primer (5'-3') | Reverse primer (5'-3') |
| --- | --- | --- |
| TSPAN17 | GAAGGGCGTTCTCTCGAACA | AAAGGCCAGGATCCCTGTTG |
| TRPA1 | TGCATGTTGCATTCCACAGAAG | TTGAGGGCTGTAAGCGGTTCATA |
| SCG5 | CTCACCAGGCCATGAATCTT | TGTTGTCTCCAGTCAACTCTGC |
| *β*-actin | GGGAAATCGTGCGTGACATTAAGG | CAGGAAGGAAGGCTGGAAGAGTG |
