## Supplementary Table 5 for "Cross-kingdom regulation of tRNAs/tRFs derived from Chinese yew"

**Supplementary Table 5. Oligonucleotides used for western blot assay**

| Oligonucleotides | Forward (5'-3') | Reverse (5'-3') |
| --- | --- | --- |
| TRPA1_siRNA01 | GGUGGGAUGUUAUUCCAUATT | UAUGGAAUAACAUCCCACCTT |
| TRPA1_siRNA02 | GAAGGACGCUCUCCACUUATT | UAAGUGGAGAGCGUCCUUCTT |
| Scramble control | UUCUCCGAACGUGUCACGUTT | ACGUGACACGUUCGGAGAATT |
